# Comparative single-nucleus RNA-seq analysis captures shared and distinct responses to beneficial and pathogenic microbes in roots

**DOI:** 10.1101/2023.08.03.551619

**Authors:** Qiuhua Yang, Zhuowen Li, Kaixiang Guan, Zhijian Liu, Ancheng Huang, Jixian Zhai, Yanping Long, Yi Song

## Abstract

Distinguishing and differentially responding to beneficial and pathogenic microbes are fundamental for plants to maintain microbiome homeostasis and promoting plant fitness. Using a recently developed protoplast-free single-nucleus RNA-seq approach, we generated single-cellular atlas of root responses to beneficial and pathogenic microbes. Notably, we identified triterpene biosynthesis as a novel cell type specific response to root pathogens and genetically confirmed the role of triterpene biosynthesis in regulating beneficial/pathogenic microbe ratios in a two-strain mixed community. Our results provide novel insights and vital resources for further elucidating novel regulators of beneficial and pathogenic microbe colonization and microbiome homeostasis.

## Main

Most eukaryotes generally live close association with complex microbial communities (i.e. the microbiome), among which beneficial, pathogenic and neutral microbes coexist^1^. Previous study suggested that plant gene can selectively regulate beneficial *Pseudomonas* colonization in the natural microbiome^2^. A recent study indicated that plant roots can distinguish a beneficial microbe from a pathogenic microbe in a two-strain community^3^. Those evidence indicates that plants can selectively shape microbiome members in a community, but how the host distinguishes and differentially responds to friends and foes and further maintaining microbiome homeostasis are poorly understood. Understanding how roots repel pathogens and recruit beneficial microbes would enable the pioneering of new strategies to maintain a healthy rhizosphere ecosystem in agriculture.

Plant roots are highly heterogeneous, and immune responsiveness varies in different zones and cell types^4,5^. To study how roots distinguish pathogens from beneficial microbes, it is fundamental to first dissect the shared and distinct responses to different microbes at the single-cell resolution. Canonical genetic and genomic approaches have substantially elucidated the immune responses to pathogens in leaves in recent decades^6,7^. Nevertheless, our understanding of plant-commensal interactions, especially in the root system, remains sparse. Bulk transcriptomic analysis has revealed tissue-specific gene expression profiles in leaves and roots upon pathogen infection^8-10^. To enhance the profiling resolution of cell type specific responses, a recent study elegantly utilized fluorescence-activated cell sorting methods to isolate different root cell types for RNA-seq^11^, which revealed more “hidden” cell type specific responses cannot be identified through bulk root RNA-seq analysis. However, a comparative analysis of cell type specific responses to pathogenic and beneficial microbes, especially at the early stage of recognition, is still lacking. The major challenge for the application of single-cell RNA-seq (scRNA-seq) in plants is that it takes several hours to do protoplast isolation after sampling. That means we cannot profile the real time transcriptional status at the sampling time point, and the enzymatic digestions will also induce the expression of stress responsive genes. Considering that plant immune responses to most elicitors can be detected within 30-90 minutes^12^, a protoplast-free scRNA-seq method can more accurately reveal the early transcriptional responses in plant-microbe interactions.

In this study, we utilized our recently developed protoplast-free single-nucleus RNA (snRNA-seq) sequencing approach to investigate root heterogeneity in shared and microbe specific response to beneficial and pathogenic microbes^13^. We chose *Pseudomonas simiae* WCS417 (WCS417 hereafter) and *Ralstonia solanacearum* GMI1000 (GMI1000 hereafter) as model beneficial and pathogenic microbes, respectively. *P. simiae* WCS417 is a model beneficial microbe identified from a naturally occurring disease suppressive soil which shows growth-promoting and disease suppressive activities^14,15^. In contrast, *R. solanacearum* GMI1000 is one of the most devasting soil-borne bacterial pathogens infecting more than 250 plant species and causing serious agricultural losses worldwide^16^. Pathogenic GMI1000 causes epidermal cell death as early as 12 hours after inoculation^4^, while beneficial WCS417 visibly promotes root hair and lateral root growth within 2 days^15^. Here we utilized a 48-well plate-based hydroponic root-microbe interaction system^2,17^ to separate roots and leaves using a mesh, and keep roots in liquid media to enable interaction with rhizosphere bacteria (Fig. 1a). To dissect whether plant roots could effectively recognize and differentially response to GMI1000 and WCS417 before visible damage or growth promoting effects occur, we performed qRT□PCR using the whole excised roots, and detected a slight upregulation of immune responsive genes (*CYP71A12, MPK11, WRKY33*) as early as 2 hours after GMI1000 treatment, and with a peak at 6 hours (Extended Data Fig. 1). In contrast, WCS417 showed a weaker induction of those immune marker genes, but higher induction of *SRO4* (AT3G47720)^18^(Extended Data Fig. 1), a WCS417 responsive gene identified from a previous RNA-seq study. This indicates that 6 hours would be an early time point with a robust transcriptional response to pathogens and beneficial microbes.

**Fig. 1.**
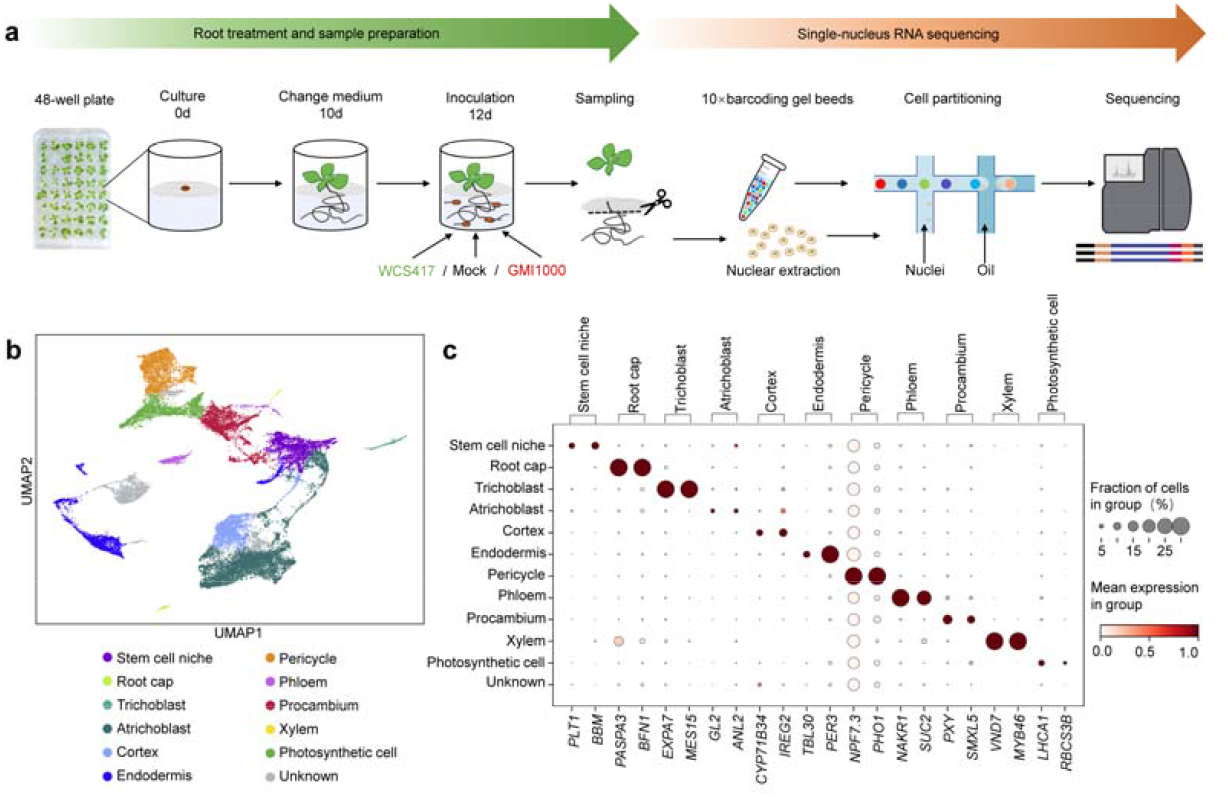
Single-nucleus RNA-seq analysis of early-stage responses to beneficial and pathogenic microbes in roots. **a**, Schematic showing the sample preparation and sequencing processes for snRNA-seq. Plants were grown in 48-well plates with liquid MS media containing 2% sucrose for 10 days before changing to fresh liquid MS media without sucrose. Plant roots were then treated with 10 mM MgSO_4_ (Mock), WCS417 or GMI1000 (final OD_600_=0.05). Samples were harvested 6 hours post treatment. **b**, UMAP visualization of 11 cell types in snRNA-seq data of *Arabidopsis* roots. **c**, Dotplot showing the expression levels of previously reported cell type-specific marker genes in 11 cell clusters.

We treated *Arabidopsis* roots with either GMI1000, WCS417 or Mock (MgSO_4_) for 6 hours, and harvested the whole roots for snRNA-seq library preparation^13^. After sequencing, data filtering and quality control, we obtained 29,214 valid nuclei from the three libraries, with 22,185 genes covering ∼80% of the genome. The median gene count and UMI count for individual cells were 751 and 990, respectively (Supplementary Table 1). We integrated three libraries by Seurat RPCAs^19^ (Extended Data Fig. 2a) and the cells were classified into 22 clusters by Leiden algorithm^20^ (Extended Data Fig. 2d). For annotation, we used CELLEX^21^ to call marker genes in each cluster (Supplementary Table 2), and analyzed their overlap with marker genes from a comprehensive *Arabidopsis* root scRNA-seq dataset recently reported by Shahan et al.^22^ (Extended Data Fig. 2b). We can assign most clusters to corresponding cell type, whereas cluster 1 and 11 showed apparently mixed cell identities. So we further divided these two clusters into subclusters and collected 26 clusters in total. Combining with cell type specific markers identified in four published root scRNA-seq datasets^23-26^, we annotated most clusters into 11 major root cell types, including stem cell niche (two clusters), root cap (one cluster), trichoblast (one cluster), atrichoblast (four clusters), cortex (two clusters), endodermis (four clusters), pericycle (two clusters), phloem (two clusters), procambium (two clusters), xylem (one cluster) and photosynthetic cells (two clusters) (Fig. 1b, Extended Data Fig.2c-d, Supplementary Table 2). The expression patterns of well-characterized marker genes also validate the accuracy of the annotation^13,22-25,27^ (Fig 1c, Supplementary Table 3).

To validate our snRNA-seq data and the treatment system, we checked the expression patterns of a few previously reported microbe-responsive genes. *Phytosulfokine receptor 1* (*PSKR1*) is a beneficial *Pseudomonas* responsive gene in roots, that is a negative regulator of root immunity and helps beneficial *Pseudomonas* bypass immunity and better colonize roots^28^. *Pseudomonas* colonization can trigger expression of a *PSKR1pro:GUS* transgene reporter in atrichoblast and cortical cells 24 hours after inoculation^28^. Our data indeed showed higher expression of *PSKR1* in diverse cell types in response to WCS417 but not to GMI1000, especially in the cortex and endodermis cells at this early stage (6 hours) of WCS417 colonization (Extended Data Fig. 3). Interestingly, we also observed that two PSKR1 ligand genes, *Phytosulfokine* (*PSK*) *1* and *2*, were both specifically induced by beneficial WCS417 but not GMI1000 (Fig. 2a, Extended Data Fig. 3). Moreover, *PSK1* and *2* showed cell type specific enriched expression in atrichoblast and phloem, respectively, which were consistent with previous reports^29^. This further supports that the PSK-PSKR pathway might be a beneficial *Pseudomonas*-specific responsive pathway in roots, and more importantly, validates the accuracy of our snRNA-seq dataset. In addition, a recent study found that WCS417 could induce a subset of suberin biosynthesis (*GPAT7*) and transporter (*ABCG16*) genes in the endodermis cells after 48 hours *Pseudomonas* colonization^11^. Our data confirmed that *GPAT7* and *ABCG16* are responsive to WCS417 and exhibited enriched expression in the endodermis cells (Fig. 2a, Extended Data Fig. 3). It is noteworthy that we not only detected consistent upregulation of suberin biosynthesis genes in response to beneficial WCS417, but also found that pathogenic GMI1000 induced an even much stronger upregulation of those genes. This highlights the importance of conducting comparative analysis of beneficial and pathogenic microbes responses in roots in order to effectively distinguish a general non-self response^30^ from a microbial lifestyle specific response.

**Fig. 2.**
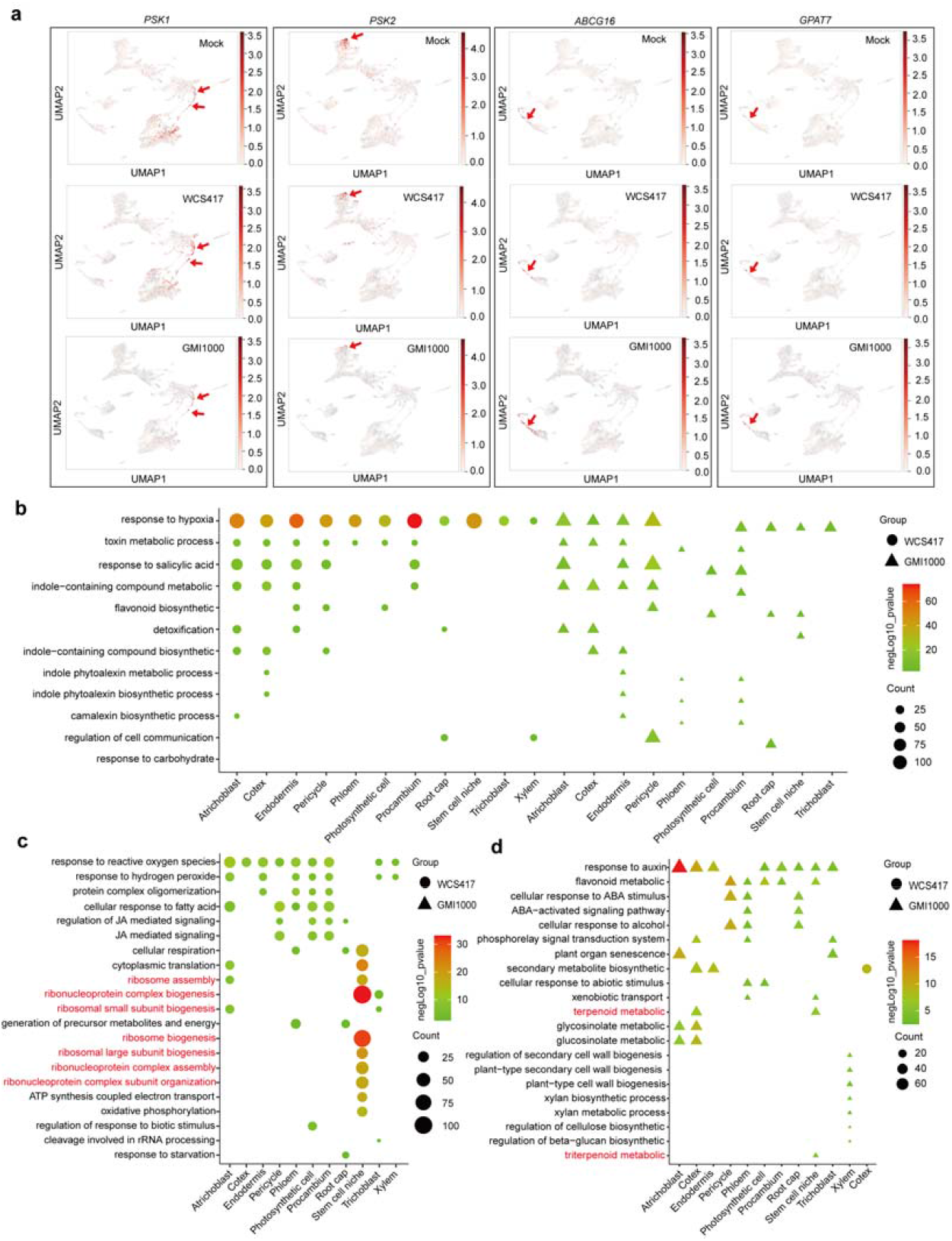
Cell type-specific responses to GMI1000 or WCS417 treatment in roots. **a**, UMAP illustration of cell type specific responses of the selected marker genes, red arrows highlight cell types in which the marker genes are up-regulated after WCS417 or GMI1000 treatment (*PSK1* in atrichoblast, *PSK2* in phloem and *ABCG16 & GPAT7* in endodermis). **b**, Shared GO terms enriched in both WCS417 and GMI1000 triggered DEGs in individual cell types. **c**, Unique GO terms exclusively or mostly only enriched in WCS417 up-regulated genes in each cell type. **d**, Unique GO terms exclusively or mostly only enriched in GMI1000 up-regulated genes in each cell type. The Y-axis represents the name of GO processes, while X-axis shows different cell types. The color of the shape indicates the negative log_10_ (P-value) of GO terms.

We next examined whether different root cell types can differentially respond to beneficial and pathogenic microbes at the transcriptome level. Both beneficial WCS417 and pathogenic GMI1000 trigger large amount of differentially expressed genes (DEGs) compared with the mock group at 6 hours after inoculation (Extended Data Fig. 4a, Supplementary table 4). However, there is very little overlap (ranging from 5% in the xylem to 18.9% in pericycle) of differentially expressed genes (DEGs) between WCS417 and GMI1000 treated roots for each cell type (Extended Data Fig. 4b). Our data strongly supports that roots can distinguish and differentially respond to beneficial and pathogenic microbes within 6 hours after inoculation, in a cell type specific manner.

To study which biological processes are involved in the response to beneficial and/or pathogenic microbes, we performed Gene Ontology (GO) enrichment analysis of WCS417 and GMI1000 induced DEGs (Fig. 2b-d). We observed diverse shared GO terms enriched in the DEGs induced by both microbes, indicating the existence of core microbial responses in roots^30^. For instance, salicylic acid (SA), phytoalexin and toxin metabolism related GO terms were highly enriched in most cell clusters upon both GMI1000 and WCS417 treatments (Fig. 2b). This is consistent with the SA related immune responses being required to avoid commensal overgrowth^28^. Intriguingly, the GO term “response to hypoxia” showed the highest enrichment in most cell clusters in response to both GMI1000 and WCS417. The strong upregulation of hypoxia related GO terms is also observed in response to immune elicitors in both roots^3^ and leaves^12^, as well as in a root autoimmue mutants^31^. Hypoxia sensing is related to the phosphorylation of calcium-dependent protein kinases and the mitogen-activated protein kinase 3/6 (MPK3/6)^32^, indicating that hypoxia signaling might be an integral part of general immune responses. Our data also indicate that the canonical innate immune responses to beneficial and pathogenic microbes share similarity, at least at this early interaction stage.

For the unique GO terms exclusively or mostly just enriched in WCS417 induced genes (Fig. 2c), we detected diverse ribosome function, translation and energy metabolism related GO terms, especially in stem cell niche (meristem), atrichoblast (nonhair) and phloem cells. This is consistent with the finding that WCS417 induce the highest (2411) and second highest (1706) amount of DEGs in stem cell niche and atrichoblast (nonhair) cells (Extended Data Fig. 4a). Ribosome function is the key bottleneck for protein translation, which is critical for growth and metabolism^33^. Moreover, several ATP synthesis- and respiration-related GO terms were highly enriched in stem cell niche (meristem). Our data successfully captured those growth promoting related cell type specific molecular features at the early stage of interaction, providing a potential mechanism underlying the well-characterized growth-promoting activity of beneficial microbes. Considering that translational regulation broadly regulates both pattern-triggered and effector triggered immunity in plants^34,35^, it would be interesting to further elucidate whether beneficial microbes induced translational reprogramming, and the significance in root-beneficial microbe interactions.

We further examined GO terms unique to GMI1000 induced DEGs, which included auxin, ABA responses and phosphorelay signal transduction related terms (Fig. 2d). We also detected that cell wall and glucan biosynthesis terms were upregulated in response to GMI1000, consistent with a previous result^36^. Notably, terpenoid biosynthesis related GO terms were exclusively enriched in GMI1000 treated roots, including cell types like stem cell niche (meristem) and cortex cells (Fig. 2d). This molecular response was not identified in a previous time-series (0-96 hours) bulk root RNA-seq dataset^36^, which might because that subtle gene expression changes of triterpenoid biosynthetic genes in a few cell types cannot be effectively detected in bulk root RNA-seq study.

Triterpenes are plant-specialized metabolites exhibiting antimicrobial activities and play essential roles in shaping the *Arabidopsis* core root microbiome^37^. Previous microbiome study identified 494 core OTUs (Operational Taxonomic Units) specifically enriched in the root microbiome of *Arabidopsis* compared with rice and wheat (both of rice and wheat do not produce triterpenes products thalianin and arabidin), while about one-third of those *Arabidopsis* specifically enriched OTUs were depleted in triterpene mutant lines (especially in *thas* mutant defective in thalianin biosynthesis)^37^. That suggests an essential role of triterpene in enriching *Arabidopsis*–specific root bacteria rather than repelling them^37^, however, in which biological context do plants produce triterpenes to shape microbiome remains unknown. Since we found that triterpene biosynthesis related genes were significantly induced by GMI1000 in several cell types, including atrichoblast, cortex and root cap cells (Fig. 3a-c), we hypothesized that roots utilize triterpenoid metabolic pathway to enrich or maintain a homeostasis of commensal colonization levels in response to pathogens. To genetically confirm this, we obtained a *thalianol synthease 1* (*thas1)* mutant, that blocked upstream of the triterpenoid (thalianol) biosynthesis and affected *Arabidopsis* microbiome structure^37^ (Fig. 3b). We then mixed WCS417 and GMI1000 (1:1 ratio) as a community to inoculate the rhizosphere Col-0 and *thas1* mutant using our 48-well plate system. We utilized antibiotic resistance to distinguish two microbes: the WCS417 strain we used has a rifampicin resistance, and we selected a natural mutant of GMI1000 that has streptomycin resistance (renamed GMI1000s). Indeed, we found that the *thas1* mutant lost the ability to control homeostasis between the colonization levels of WCS417 and GMI1000s. As evidenced by significantly lower WCS417 levels and WCS417/GMI1000s ratios (Fig. 3d-f). Our data revealed a terpenoid biosynthetic pathway selectively enrich beneficial *Pseudomonas* in a community, and provides a validation of using our snRNA-seq dataset to discover novel genes/pathways that regulate commensal colonization.

**Fig. 3.**
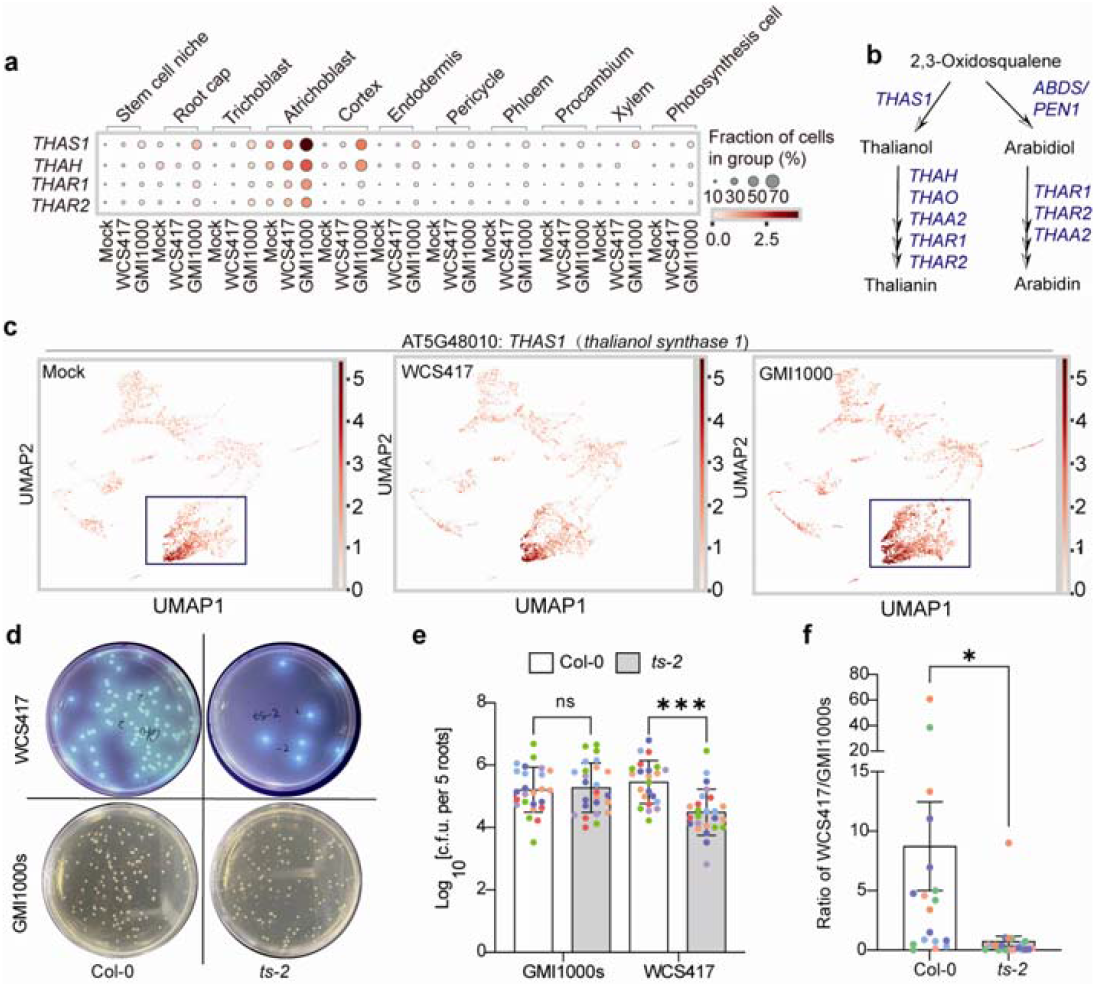
The triterpenoid biosynthetic pathway is required to selectively regulate WCS417 colonization in a WCS417-GMI1000 mixed community. **a**, Several triterpenoid biosynthetic genes are strongly induced by GMI1000 but not WCS417. **b**, A simplified pathway shows the key enzymes catabolizing the biosynthesis of two triterpenoid products (thalianin and arabidin). **c**, UMAP illustration of the expression patterns of the *THAS1* gene in mock- and microbe-treated roots. **d**, Representative images of WCS417 plates and GMI1000s plates from Col-0 and *ts-2*. **e**, The colonization levels of WCS417 and GMI1000s in Col-0 and the*ts-2*mutant. Data are from 5 independent experiments, are different colors indicate data points from different individual experiments. The two strains were co-inoculated in the rhizosphere in a 48 well plate, and CFUs were measured 60 hours after inoculation. **f**, Ratios of CFU (WCS417)/CFU (GMI1000s) from the roots of Col-0 and the *ts-2* mutant.

How eukaryotic organisms engage with beneficial microbes while restricting pathogenic microbes in complex microbiomes is a fundamental question in the host-microbiome interaction field. To study this, it is essential to first know whether and how roots can distinguish between beneficial and pathogenic microbes. In this work, we found that all root cell types could differentially respond to “friend and foe” in roots even within the first 6 hours of interaction. Our snRNA-seq data and genetic confirmation further provided novel insights that specific secondary metabolites, such as triterpenes could modulate beneficial/pathogenic microbe homeostasis in a community. Considering the broad interest in root immune responses as well as how roots distinguish beneficial and pathogenic microbes, we have provided a website (https://zhailab.bio.sustech.edu.cn/sn_microbe) for the community to further explore gene expression patterns at a single-cell resolution. We believe our insights from those two model strain systems will provide critical clues for further identifying plant genes shaping root associated microbiomes and maintaining beneficial/pathogenic microbe homeostasis.

## Methods

### Plant material and growth condition

*Arabidopsis* seeds were surface sterilized with 20% bleach for 1-2 min, followed by 75% ethanol for another 1-2 min, and then washed three times with sterilized water. Afterwards, seeds were imbibed at 4 □ in the dark for 4 days. Plants were grown in growth rooms at 23°C under 16h light/8h dark period with light intensity at 100□ μmol□ m^−2^ s^−1^. For the 48-well plate assay, plants were grown in 48-well plates with liquid 1×Murashige and Skoog (MS) medium containing 2% (w/v) sucrose for 10 days and transferred to fresh liquid 1×MS medium without sucrose for 2 more days before inoculation.

### Bacterial inoculation and sampling

For snRNAseq and qRT-PCR experiments, *Arabidopsis* seedlings were grown in 48-well plates with liquid 1× MS medium containing 2% (w/v) sucrose for 10 days, and then the media was changed to fresh liquid MS medium without sucrose for 2 more days. For bacterial inoculation, *P. fluorescens* WCS417 and *Ralstonia solanacearum* GMI1000 were washed twice and resuspended in 10 mM MgSO_4_ to remove potential exudated elicitors during overnight culture. WCS417 and GMI1000 were diluted to OD_600_ = 0.5 in 10 mM MgSO_4_ and then added to the liquid rhizosphere to a final OD_600_ of 0.05. Liquid MS medium supplemented with the equal volume of 10 mM MgSO_4_ was used as a mock control. For qRT-PCR the roots were harvested at 2 h, 6 h, 24 h, 30 h, 48 h after inoculation and about 5-10 roots were pooled as one sample. For snRNA-seq roots were pooled from 30-40 seedlings at 6 hours after inoculation as one sample. After harvest, the roots were snap-frozen in liquid nitrogen, and stored at -80 □.

### Nuclei isolation and 10× snRNA-seq library construction

The roots were chopped thoroughly in ice-cold nuclei isolation buffer (1× NIB buffer (Sigma-Aldrich, #CELLYTPN1), supplemented with 1 mM dithiothreitol (DTT, Thermo Fisher, R0861), 1× protease inhibitor (Sigma, 4693132001) and 0.4 U/μL murine RNase inhibitor (Vazyme, R301-03)). Then the lysate was passed through a 40 μm strainer (Sigma, CLS431750-50EA) and centrifuged at 500 × g for 5 min at 4 °C. The pellet was gently resuspended in 500 μL of nuclei isolation buffer and 0.5 μL of 1 mg/mL DAPI was added. To remove the remaining tissue debris, the nuclei suspension was filtered through a 20 μm cell strainer (pluriStrainer Mini, 43-10020). For cell sorting, the nuclei were loaded to flow cytometer (Sony MA900) equipped with a 100 μm nozzle. A total of 100,000 nuclei for each sample were sorted into a 15 mL tube with 2 mL collection buffer (1× PBS, supplemented with 1% BSA and 0.4 U/μL murine RNase inhibitor). The nuclei were centrifuged at 500 × g for 5 min at 4 °C and resuspended in 50 μL collection buffer. The quality and yield of nuclei were checked under a fluorescence microscope using the DAPI channel. About 16000 nuclei of each sample were loaded into the 10x Genomics chip for cell barcoding. The libraries were prepared following the manufacturers’ instructions with 10x Chromium Single Cell 3’ Solution v3.1 kit. The pair-end libraries were sequenced with Illumina Nova-seq instrument.

### RNA extraction and qRT-PCR

For qRT-PCR assay, total RNA was extracted with the FastPure Plant total RNA isolation Kit (Vazyme 7E602I2) and quantified by nanodrop. Approximately 1 μg of RNA was reverse-transcribed to cDNA with HiScript III All-in-one RT SuperMix (Vazyme Q711-03). The qPCR was performed with ChamQ Universal SYBR qPCR Master Mix (Vazyme R333-01) with the housekeeping gene EIF4A as an internal control. The primer pairs used for qRT-PCR were listed in Supplementary Table 5.

### Data processing and cell type annotation

The raw reads were preprocessed by Cell Ranger (v6.0.0)^38^ and aligned to *Arabidopsis* TAIR 10 genome, generating the h5ad files for each sample. Then we used SCANPY package (v1.8.0)^20^ in python to read the files and used ScDblFinder (v1.10.0)^20^ to remove the doublet. After that, we filtered out the genes that detected in less than 10 cells, and kept only the cells with gene count larger than 300 and less than 3500, and UMI count larger than 500 and less than 6000. In addition, cells with more than 0.05% transcripts from mitochondria or chloroplasts were removed.

We then adopted Seurat (v. 4.3.0)^19^ to integrate the three matrixes following the official guidance (https://satijalab.org/seurat/articles/integration_rpca.html). We used SCTransform to normalize and called top 3000 highly variable features for each matrix, and selected the shared high variable features for canonical correlation analysis (CCA) to identify the integrating anchors. After that we used the ‘scanpy.pp.neighbors’ function to call the neartest-neighbour graph (default parameter), and performed leiden alogorithm (scanpy.tl.leiden, resolution = 0.6) on the graph for clustering. The Uniform Manifold Approximation and Projection (UMAP) algorithm was used for visualization (scanpy.tl.umap, resolution = 0.2). For annotation, we first used CELLEX (CELL-type Expression-specificity, v1.2.2^21^) to call the cluster-enriched genes from our data and the reference data (Shahan et al., 2022)^22^, and annotated the clusters according to their Intersection of Union level (Extended Data Fig. 2c).

### Differential expression analysis and GO enrichment analysis

We adopted Diffxpy (https://github.com/theislab/diffxpy) to identify the Differential Expressed Genes (DEGs) from the treatment of GMI1000 or WCS417 against the buffer-treat sample within each cluster. Genes expressed in less than three cells in the two comparing clusters would be masked for DEG calling. From the Diffxpy calculation result, genes with p-value < 0.05 and log2 (fold change) > 1 would be considered significantly upregulated (UP); and p-value < 0.05 and log2 (fold change) < -1 would be considered significantly downregulated (DOWN). We used the enricher function in R package clusterProfiler (v4.6.0)^39^ for GO enrichment analysis of the DEGs, and the dotplot illustration of GO results was conducted by the ImageGP website^40^.

### Quantifying microbe colonization levels in roots

Seedlings grown in our 48 well plate system were used for the root colonization assays. A mixture of two microbes (WCS417 and GMI1000) at the 1:1 ratio was added to a final concentration of OD=0.00002 at day 12. After 48 hours inoculation, we harvested roots and pooled 5 roots as one sample for series dilution and CFU calculation. The WCS417 strain we used has a rifampicin resistance, and we used rifampicin plates to selective quantify WCS417 in the two-stain community. To distinguish GMI1000, we streaked 5 ml overnight culture of GMI1000 on 10 streptomycin (50 mg/ml) plates, and we were able to select a streptomycin resistant GMI1000 strain (GMI1000s). The WCS417 plates pictures were taken under UV lights, which enable us to show the fluorescence of WCS417 colonies.

## Data availability

The snRNA-seq data will be deposited in the China National Center for Bioinformation database upon publication.

## Code availability

All the codes used in this work could be found at Zhai lab’s github site: https://github.com/ZhaiLab-SUSTech/snRNA-seq_microbes

## Acknowledgements

We thank Dr. Cara Haney and Dr. David Thoms for critical reading of the manuscript. The group of Y.S. was supported by the NSFC General Project (32270286) and the Stable Support Plan Program of Shenzhen Natural Science Fund Grant (20220815160107001). The group of J.Z. was supported by the National Key R&D Program of China Grant (2019YFA0903903); the Program for Guangdong Introducing Innovative and Entrepreneurial Teams (2016ZT06S172); the Shenzhen Sci-Tech Fund (KYTDPT20181011104005); the Key Laboratory of Molecular Design for Plant Cell Factory of Guangdong Higher Education Institutes (2019KSYS006); the Stable Support Plan Program of Shenzhen Natural Science Fund Grant (20200925153345004); and the Center for Computational Science and Engineering at Southern University of Science and Technology.

## Author information

Y.S., Y.L., J.Z. and A.H. designed the project. Q.Y., Y.L., K.G. and Y.S. performed the experiments. ZW.L., K.G., Q.Y., and ZJ.L. analyzed the data. Y.S., Q.Y., ZW.L. and Y.L. wrote the manuscript.

**Extended data Fig.1.**
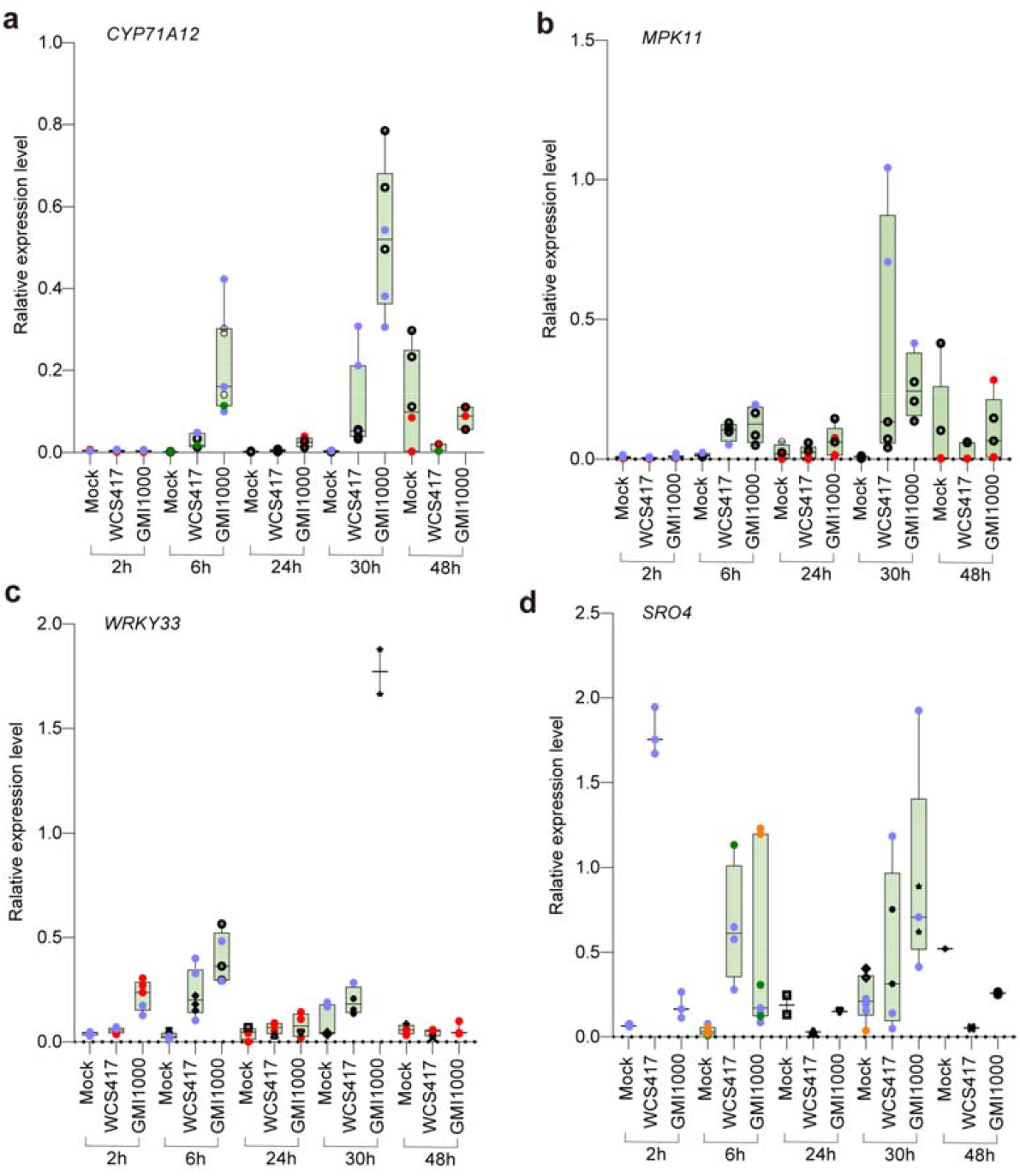
Time-series detection of the pathogen and beneficial microbe responsive genes in roots by qRT-PCR. **a-c**, Immune related *CYP71A12, MPK11* and *WRK33* showed stronger responsiveness to GMI1000, while *SRO4* (**d**) showed high responsiveness to WCS417. Dots in different colors indicate biological replicates from independent experiments. The expression of *EIF4A* gene was used as an internal control.

**Extended data Fig.2.**
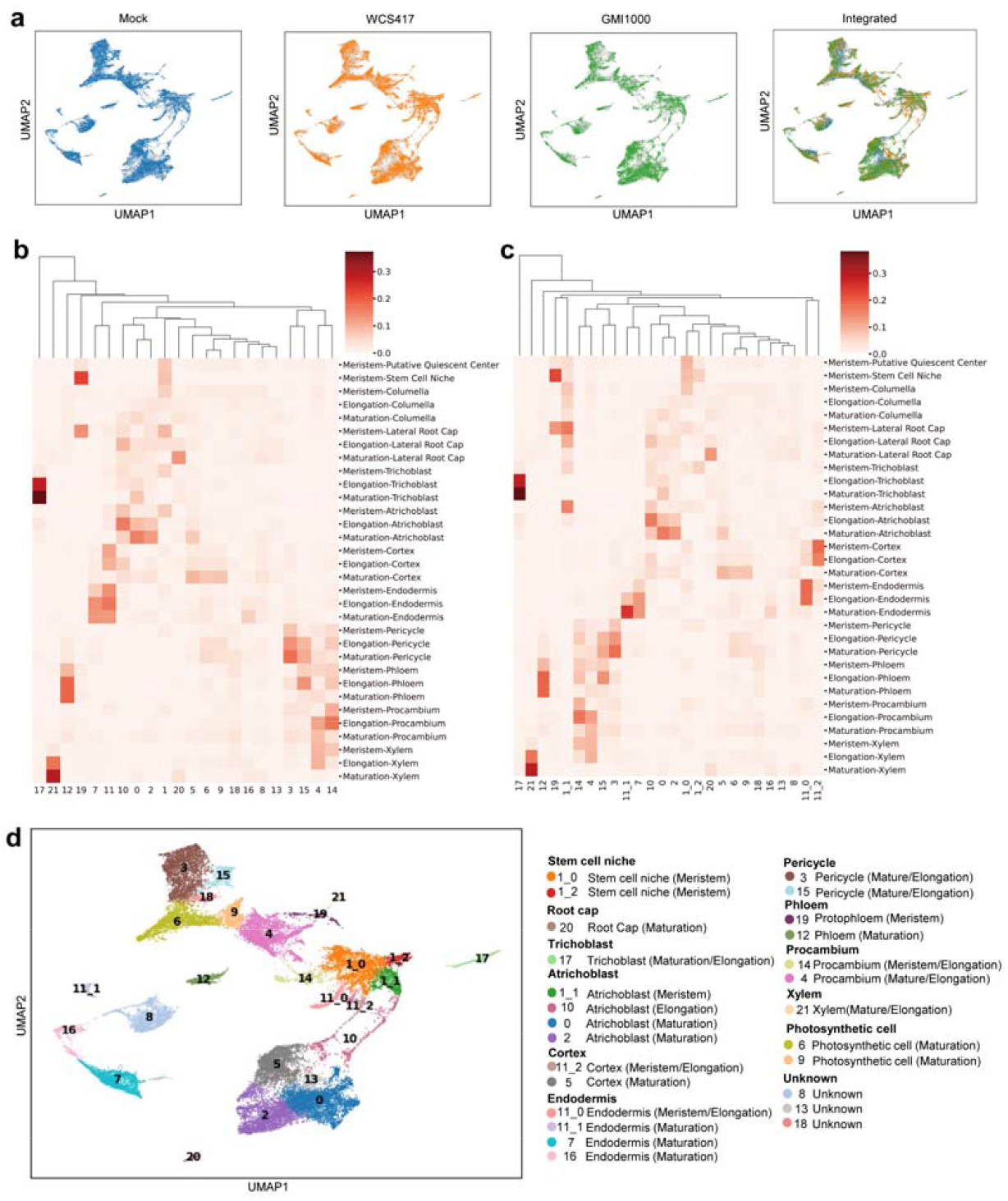
Integration and annotation of the snRNA-seq data. **a**, Visualization of the three snRNA-seq libraries and their integrated results with UMAP. **b**, Intersection of Union (IOU) of the cluster marker genes from our snRNA-seq atlas (leiden resolution = 0.6) and from a comprehensive *Arabidopsis* root scRNA-seq atlas reported by Shahan *et al*.,^24^. The cluster marker genes were called with CELLEX (CELL-type EXpression-specificity, v1.2.2^21^), and keep the genes with enrich score > 0.8. **c**, Intersection of Union (IOU) of the cluster marker genes from our snRNA-seq atlas with sub-clusters and from the well-annotated atlas reported by Shahan *et al*.,^24^. Subclusters from cluster 1 (named 1_0, 1_2, 1_2) and from cluster 11 (named 11_0, 11_2, 11_2) were added for marker calling. **d**, Annotation results for all 26 clusters in UMAP.

**Extended data Fig.3.**
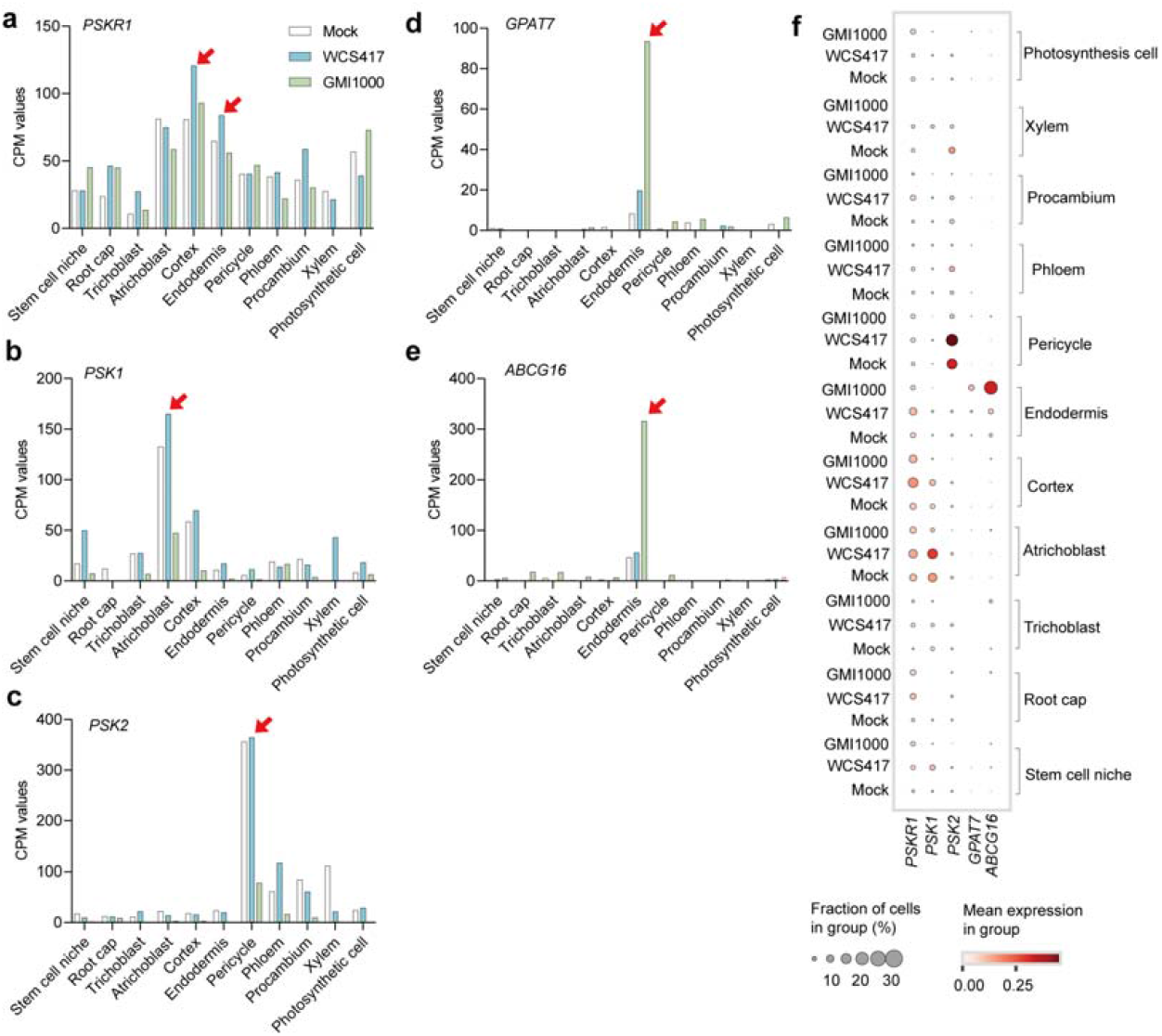
The expression patterns of the *PSK-PSKR1* pathway and suberin biosynthesis genes. **a-e**, The CPM (counts per Million) values of each gene in different cell clusters in Mock and microbe treated roots, red arrows highlight genes up-regulated in different cell types. **f**, Dotplot illustration of the expression patterns of different genes in each cluster. The fraction of cells expressing each gene were reflected in dotsize, and the expressing levels were indicated by color.

**Extended data Fig.4.**
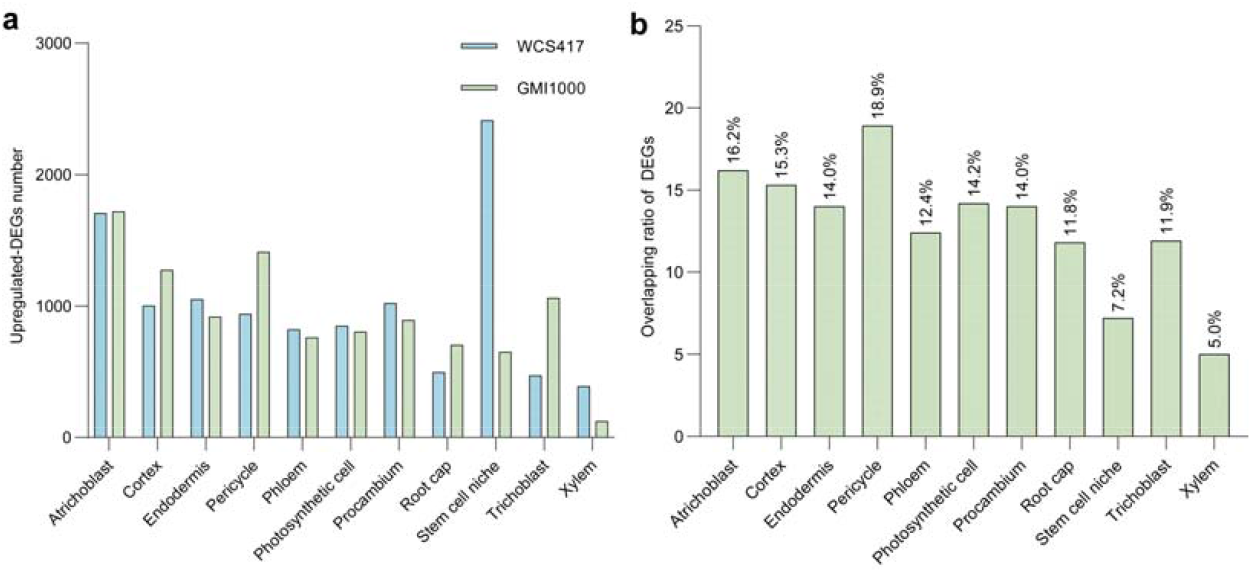
The number and overlapping ratios of DEGs after GMI1000 or WCS417 treatment in each cell type. **a**, The number of up-regulated DEGs in all 9 root cell types after WCS417 or GMI1000 treatment, all DEGs were compared with the mock treated group in each cell type; **b**, the overlapping ratios between WCS417- and GMI1000 induced DEGs in each cell type. The ratios were calculated by using the number of shared DEGs relative to the total DEGs numbers with each cell type.

